# Programmed withdrawal of cilia maintenance followed by centriole capping leads to permanent cilia loss during cerebellar granule cell neurogenesis

**DOI:** 10.1101/2023.12.07.565993

**Authors:** Sandii Constable, Carolyn M. Ott, Andrew L. Lemire, Kevin White, Amin Lim, Jennifer Lippincott-Schwartz, Saikat Mukhopadhyay

## Abstract

Primary cilia in brain neurons provide a privileged compartment for binding and responding to extracellular ligands such as sonic hedgehog. Paradoxically, cilia in differentiating cerebellar granule cells are deconstructed during neurogenesis. To identify mechanisms underlying this newly defined cilia deconstruction pathway, we used single cell transcriptomic and immunocytological analyses to compare the transcript and protein signatures of differentiating and progenitor granule cells. We found that differentiating granule cells lacked transcripts for key regulators of pre-mitotic cilia resorption, suggesting cilia disassembly in differentiating cells was distinct from pre-mitotic cilia resorption. Further analysis revealed that during differentiation, transcription of genes required for cilia maintenance decreased. Specifically, protein components of intraflagellar transport complexes, pericentrosomal material and centriolar satellites all decreased as granule cells matured. The changes in transcription and translation correlated with the downregulation of sonic hedgehog signaling at the onset of differentiation. We also found binding of centriolar cap proteins to the mother centrioles as granule cell neurons matured. These data indicate that global, developmentally programmed, diminution of cilium maintenance caused cilia deconstruction in differentiating granule cells. Furthermore, the capping of docked mother centrioles prevents cilia regrowth likely blocking dysregulated sonic hedgehog signaling and tumorigenesis.

## INTRODUCTION

Primary cilia are microtubule-based structures that are templated from mother centrioles and function as signaling hubs in diverse cellular contexts (Anvarian et al., 2019; Kumar and Reiter, 2021; Mill et al., 2023; Nachury and Mick, 2019). In the accompanying paper we established that granule cell (GC) neurons in the cerebellum lack cilia (Ott, Constable et al. co-submitted). Early during GC neuron differentiation, the cilia are disassembled in a process we call cilia deconstruction. Once lost, the cilia do not regrow despite docking of the mother centrioles at the plasma membrane (Ott, Constable et al. co-submitted). The deconstruction of GC neuronal cilia is remarkable because neurons in most other brain regions are ciliated (Green and Mykytyn, 2014; Lee and Gleeson, 2011; Louvi and Grove, 2011; Ott et al., 2023; Wu et al., 2023). Aberrant growth of GCs causes medulloblastoma, the most common malignant pediatric brain tumor (Northcott et al., 2011). Given that subtypes of medulloblastoma cells can regain cilia and responsiveness to the mitogen sonic hedgehog (SHH) similar to the GC progenitors that give rise to GC neurons (Chizhikov et al., 2007; Han et al., 2009; Spassky et al., 2008; Youn et al., 2022), cilia deconstruction could be critical for preventing tumor growth and in designing therapeutics.

The regulation of cilia disassembly has previously been studied in cultured cells prior to cell division (Pugacheva et al., 2007; Tucker et al., 1979). However, cilia deconstruction during GC neurogenesis occurs in post-mitotic differentiating cells, so the known pathways of ciliary disassembly might not be relevant. Also, in contrast to cultured cells, GC cilia deconstruction occurs is a complex multicellular environment in the developing cerebellum. GC progenitors proliferate in response to coincidental inputs from SHH, and laminin present in the extracellular matrix close to the pia (Ong et al., 2020). Upon differentiation, dramatic reprogramming propels the GCs along glial projections away from the pia (Leto et al., 2016). Completion of GC neuronal maturation upon arrival at the internal granule layer (IGL) involves dendritic maturation and establishment of synaptic connections (Leto et al., 2016). Progenitor cells in the outer external granule layer (EGL) all have cilia (Chizhikov et al., 2007; Spassky et al., 2008), which are resorbed prior to mitosis and then reassembled after mitosis (Ott, Constable et al. co-submitted). Once these progenitor cells stop dividing and begin to differentiate, cilia are deconstructed either before or as GCs migrate from the inner EGL through the molecular layer (ML) to the IGL (Ott, Constable et al. co-submitted) (Figure 1A). It is unclear what causes primary cilia deconstruction in differentiating GCs, whether this process shares any features with the well-known pre-mitotic cilia resorption pathway in proliferating cells (Liang et al., 2016; Malicki and Johnson, 2017; Wang and Dynlacht, 2018) and how cilia reassembly is prevented from docked centrioles in mature GC neurons.

**Figure 1.**
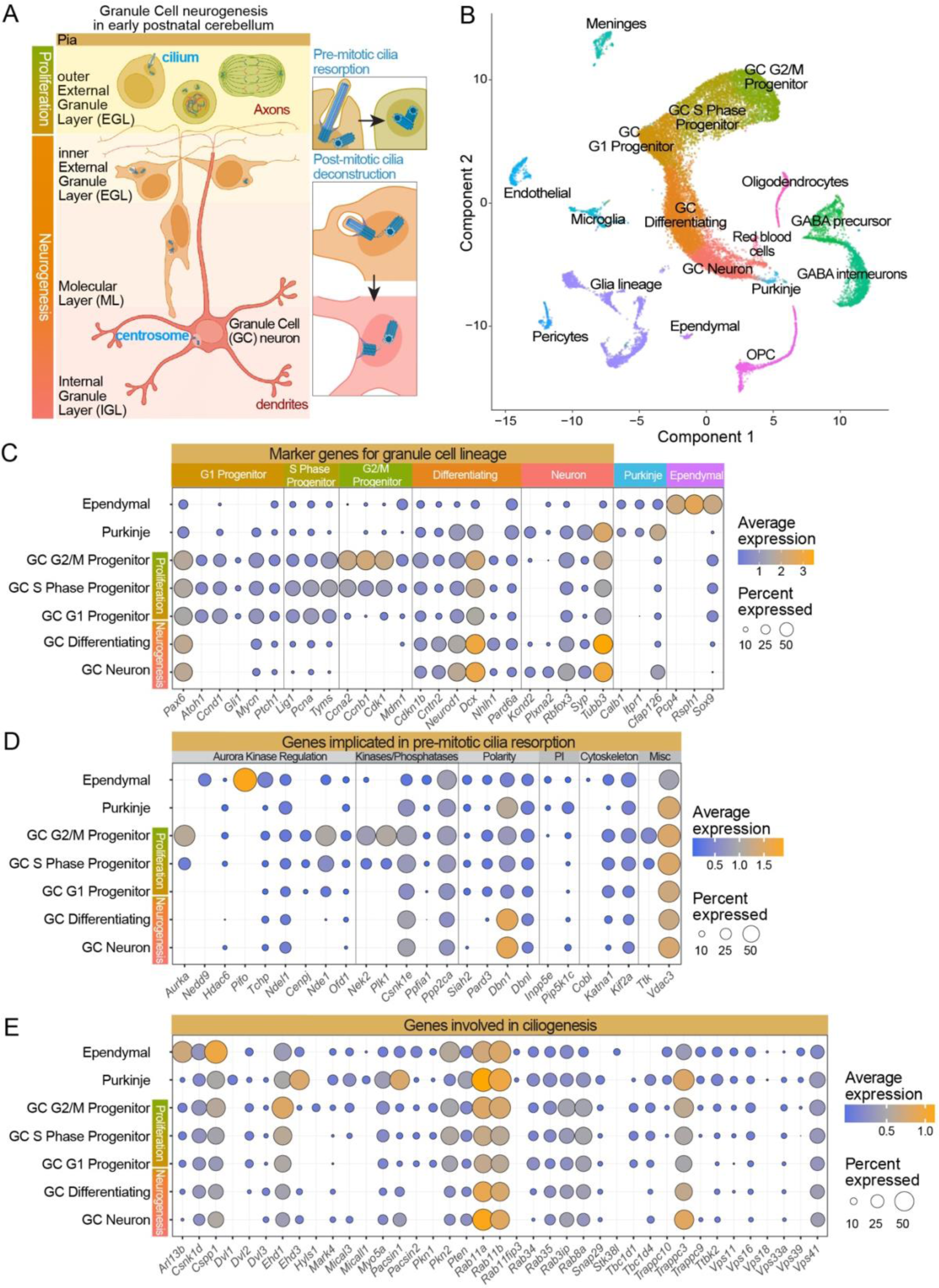
Transcriptomic differences between proliferating and differentiating GCs suggest unique regulation of cilia deconstruction is post-mitotic cells. (A) During neurogenesis, progenitors GCs proliferate in the outer EGL. Differentiating GCs, which migrate as they mature, begin in the inner EGL and move toward the IGL. GCs in the outer EGL resorb cilia prior to mitosis. Differentiating GCs begin with small internal, concealed cilia that are deconstructed during neuronal maturation. The mother centriole of mature GCs dock at the plasma membrane without extending a cilium (Ott, Constable, co-submitted). (B) scRNA-seq datasets from P5, P7 and P14 mice (Vladoiu et al., 2019) were combined, and cells were clustered based on differential expression. Clusters are plotted as a UMAP, and the identity of different cell types are labeled. (C) Expression of representative identifying genes expressed in the GC lineage were plotted. Representative genes for Purkinje neurons and ependymal cells are also included. (E and F) Expression of genes known to be involved in pre-mitotic cilia disassembly, severing or ciliogenesis were plotted for each cluster in the GC lineage. Ciliated Purkinje neurons and ependymal cells were included for comparison. The color of the dot represents the average expression level across cells and the size of the dot is determined by the number of cells in which transcription of the gene was detected.

To identify the molecular mechanisms that accomplish cilia deconstruction and prevent reciliation in differentiating GCs we used single cell transcript analysis and immunocytochemistry. These strategies captured developmental processes in the native cerebellum and leveraged the transcriptional reprogramming that occurs upon onset of neurogenesis. We discovered that many cilia and centrosome transcripts and proteins diminished as GC differentiation progressed. We determined that unlike pre-mitotic cilia resorption, cilia deconstruction in differentiating GCs was not a linear signaling cascade, but rather programmed diminution of proteins supporting cilium and centrosome maintenance. In addition, we found cilia regrowth was prevented by capping of the mother centrioles as GC neurons matured. We show that regulated cilia deconstruction contributes to loss of SHH responsiveness during GC differentiation. Although it is known that loss of individual cilia or centrosome proteins can cause pathological cilia disassembly, this is the first evidence that global reductions in cilia and centrosome proteins during differentiation removes cilia.

## RESULTS

### Mechanisms of ciliary deconstruction in differentiating GCs are distinct from pre-mitotic ciliary disassembly

Key mediators for pre-mitotic cilia resorption have already been identified in cultured cells (Liang et al., 2016; Malicki and Johnson, 2017; Wang and Dynlacht, 2018), so we investigated whether the same regulatory mechanisms cause cilia deconstruction in post-mitotic, differentiating GCs. Because dramatic transcriptional reprogramming occurs when GC progenitors commit to differentiation (Ha et al., 2019; Sato et al., 2008), we compared the transcript levels of known cilia disassembly regulators in progenitor and differentiating GCs by utilizing previously published single cell transcriptomic data of developing mouse cerebella (Vladoiu et al., 2019). We pooled the P5, P7, and P14 datasets, which include proliferating and differentiating GCs (Leto et al., 2016) Cells were clustered by similarity and assigned identities based on expression of established marker genes (Figure 1B, Sup. Figure 1, and Sup. Table 1). In this UMAP (Uniform Manifold Approximation and Projections) plot, the cluster of GC progenitor and mature cells were a continuum which were separated into five clusters based on cell marker expression. G1 progenitors can either divide or begin to differentiate (Figure 1B). G1 progenitors were found in the center of the plot with cell cycle progression clusters extending above, and differentiation and maturation clusters extending below. The expression of representative identifying transcripts for each GC cluster, as well as for Purkinje neurons and ciliated ependymal cells, are presented in Figure 1C.

To investigate similarities in cilia disassembly, we compared the expression of known pre-mitotic cilia disassembly promoting genes between the S and G2/M phase GC progenitors and differentiating/mature GCs (Figure 1D, Sup. Table 2). As expected, genes encoding proteins known to promote cilia resorption were detected in the S and G2/M clusters, some at high frequency (Figure 1E). However, we found that transcripts essential for triggering pre-mitotic cilia resorption were downregulated in differentiating and mature GCs during post-mitotic cilia deconstruction. An important example is Aurora kinase A (AURKA) which functions as a nexus for a multitude of signals in promoting cilia disassembly (Pan et al., 2004; Pugacheva et al., 2007). *Aurka* expression was detected at high levels in a large percent of G2/M progenitors but not in differentiating or mature GCs. We also examined transcription of factors that interact with or are regulated by AURKA (Kinzel et al., 2010; Liang et al., 2016; Malicki and Johnson, 2017; Wang and Dynlacht, 2018). Like *Aurka*, expression levels of *Hef1* (*Nedd9*), *Hdac6*, *Pifo* (*Pitchfork*), and *Tchp* (*trichoplein*) were either undetected or did not change dramatically. CENPJ (also known as CPAP), functions as a scaffold in recruitment of a cilium disassembly complex that consists of AURKA, NDE1 and OFD1, to the ciliary base in neuron progenitors (Gabriel et al., 2016). Like *Aurka*, *Cenpj, Nde1, and Ofd1* were expressed at high levels in S and G2/M progenitors but not in differentiating and mature GCs.

Additional kinases impact pre-mitotic ciliary resorption including the NIMA-related kinase, NEK2 (Kim et al., 2015), the polo-like kinase 1 (PLK1) (Wang et al., 2013), and TTK (Doornbos and Roepman, 2021). These kinases are highly expressed in the S and G2/M phase GC clusters and undetected in the differentiating and mature GCs. The microtubule depolymerizing kinesin KIF2A, a target of PLK1 (Miyamoto et al., 2015; Piao et al., 2009), was also detected with less frequency in differentiating and mature GCs but was not downregulated to the same extent as *Plk1*.

A small group of genes involved in cell polarity have also been implicated in pre-mitotic cilia resorption (Nager et al., 2017; Ong et al., 2020). Two of these, *Siah2* and *Pard3* are expressed at higher levels in proliferating GC progenitors. In contrast, transcripts of the actin binding protein *Debrin (Dbn1)* increases and *Debrin-like (Dbnl)* is maintained during GC neuron maturation, perhaps because these gene products participate in nucleokinesis and migration (Trivedi et al., 2017).

Another mechanism for cilia removal in response to stress, pharmacological induction or serum addition is rapid shedding of cilia (Liang et al., 2016; Mirvis et al., 2019). To investigate whether severing could be the mechanism for cilia loss during differentiation, we assessed the transcript levels of katanin, the microtubule severing enzyme implicated in cilia severing (Esparza et al., 2013; Lohret et al., 1998; Mirvis et al., 2019). We found *Katna1* levels to be diminished in differentiating and mature GCs. This lack of evidence for severing is consistent with data in our accompanying paper where we found only a single cilium with constriction suggestive of cilia severing, in contrast to the hundreds of examples of shortened cilia that favors a cilia deconstruction process (Ott, Constable et al, co-submitted). Because ciliary membrane loss can occur through release of membrane vesicles (Cao et al., 2015; Nager et al., 2017; Phua et al., 2017; Wang and Barr, 2016; Wood and Rosenbaum, 2015) we also examined the levels of *Inpp5e*, a phosphoinositide 5’phosphatase involved in the decapitation of ciliary tips (Nager et al., 2017; Phua et al., 2017). *Inpp5e* transcripts were only detected in G2/M GCs, not in differentiating or mature neurons.

In summary, transcripts for proteins that promote cilia resorption were largely reduced or undetected in differentiating and mature GCs, suggesting that the reported mechanisms of pre-mitotic cilia disassembly, while active in cycling GC progenitors, were likely not primarily utilized during cilia deconstruction in post-mitotic GC neurons.

### Ciliogenesis promoting genes are not globally downregulated during cilia deconstruction

To investigate alternative mechanisms for cilia disassembly in differentiating GCs, we first looked to see if there was simply a decrease in transcription of ciliogenesis promoting genes. We compared the expression of an extensive collection of ciliogenesis promoting genes between the GC subtypes (Figure 1F, Sup. Table 1). The transcript levels in ciliated Purkinje neurons and ependymal cells were included in the analysis for comparison. We found no global downregulation in differentiating and mature GCs, although some pivotal ciliogenesis genes, including *Hyls1*, *Hhd1*, and *Rab8a* (Zhao et al., 2022) and *Rab34*, which regulates intracellular ciliogenesis (Ganga et al., 2021; Stuck et al., 2021), were decreased during differentiation. Several ciliogenesis genes were transcribed at higher levels in differentiating and mature GC neurons compared to cycling GC progenitors, including *Dvl3, Ehd3, Pacsin1, Rab11a, Rab11b,* and *Trapp II* complex genes (Zhao et al., 2022). Some of these genes, including *Rab11a/b*, and the TRAP II complex also contribute to cell polarization, the secretory pathway and membrane trafficking in multiple contexts (Jing and Prekeris, 2009; Kim et al., 2016). Another gene involved in ciliogenesis with higher levels of transcript in mature neurons was *Pacsin1.* PACSIN1 has been implicated in formation of membrane tubules from ciliary vesicles (Insinna et al., 2019), features that we notice during ciliary deconstruction in differentiating GCs (Ott, Constable et al. submitted). We conclude that although transcriptional regulation of individual core ciliogenesis program genes could impact cilia deconstruction, evidence for a mechanism based on collective downregulation is lacking in our data.

### Expression pattern clustering revealed global decreases in many cilia and centrosome genes

Extensive transcriptional changes accompany both the onset and completion of GC differentiation (Ha et al., 2019; Sato et al., 2008), so we wondered if genes that stimulate cilia deconstruction during this process could be upregulated. To identify potential candidates, we grouped genes based on normalized expression patterns across the GC subpopulations. To enhance our chances of finding relevant factors, we pooled the top 5% of detected transcripts with a curated list of cilia and centrosome genes (Sup. Table 2). We used a cutoff value (k value) of 10 in the dendrogram to identify gene expression groups (Sup. Table 3) and then performed gene ontology analysis using DAVID (Sherman et al., 2022) to determine pathway components enriched in each expression cluster (Sup. Table 4). Figure 2A shows the dendrogram and the normalized expression pattern of all genes in each group. The expression patterns and DAVID analysis were like those generated using the top 5% of transcripts alone and a k value of 9 (Sup. Table 5 and 6). Genes were not distributed equally across the expression groups: approximately 72% of cilia genes from the curated list were in groups 1, 5 and 2, whereas the centrosome genes were distributed across groups 1, 2, 5, 7, and 9 (Figure 2B).

**Figure 2.**
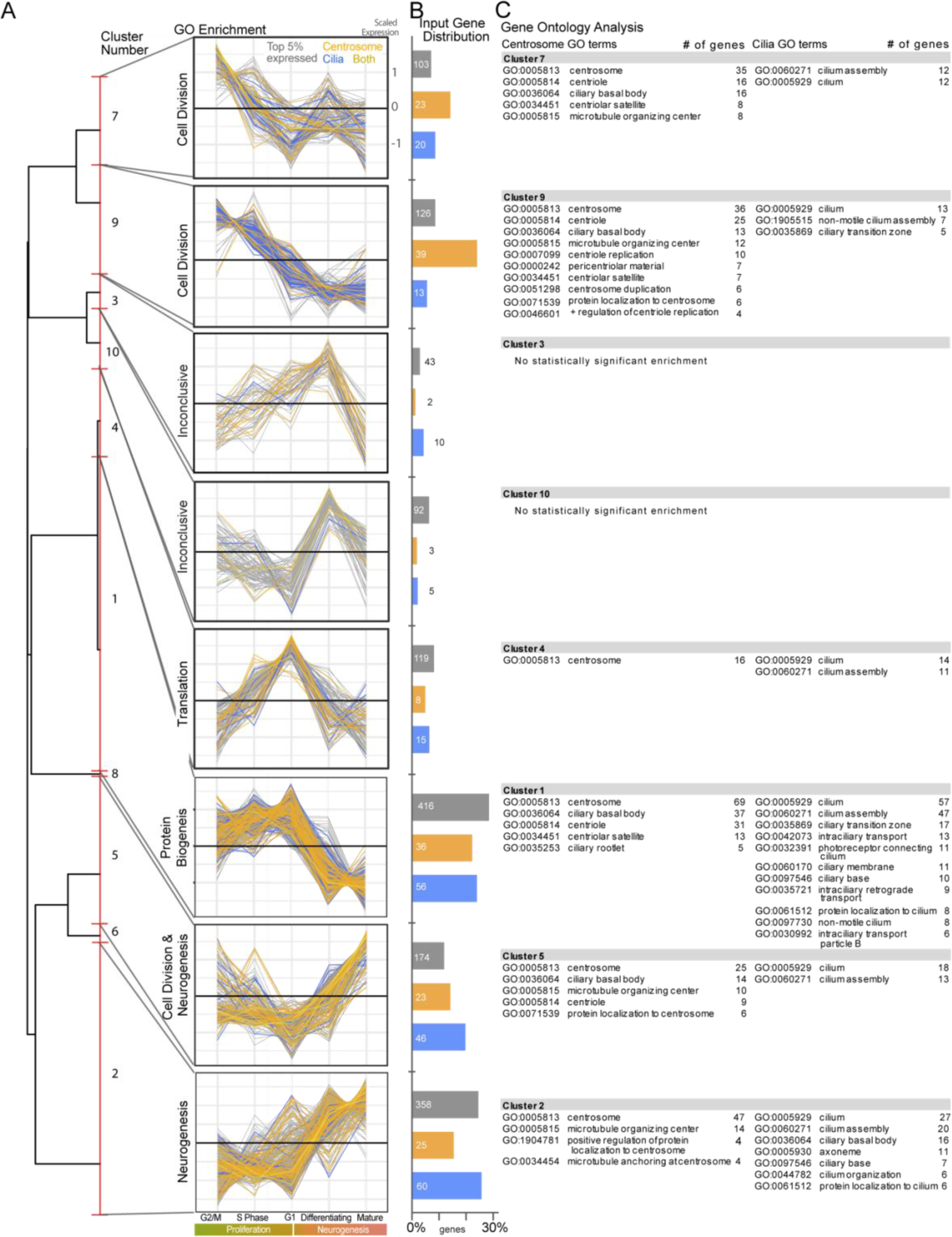
Clustering of genes by expression patterns reveals changes in cilia and centrosome genes during GC differentiation. (A) The expression of the top 5% of detected transcripts combined with a curated list of cilia and centrosome genes was normalized and then genes were clustered based on expression pattern across the GC lineage. The dendrogram of clusters is shown at the far left. The red line represents cutoff value that created the 10 clusters. The clusters were analyzed using DAVID (Sherman et al., 2022) and similar, highly significant GO terms were used to label the clusters where GO terms were significantly enriched. In the graphs of the normalized expression pattern in each cluster, the genes from the top 5% are grey, centrosome genes are orange, cilia genes are blue and genes that are both cilia and centrosome proteins are yellow. (B) The distribution of input genes in each cluster is graphed as a percentage of the total for the top 5%, centrosome or cilia genes. The number of genes is also included for each bar on the graph. (C) The significantly enriched centrosome and cilia related GO terms are listed for each cluster. The number of genes in the cluster with each GO term is indicated.

Before looking specifically at the cilia and centrosome genes, we sought to understand more about the genes in each cluster. We identified statistically enriched GO terms within the expression pattern groups. Predominant themes in the GO terms provided insights into the changes in biological pathways. Specifically, group 1 included several GO terms related to protein biogenesis, groups 2 and 5 included those related to neurogenesis, group 4 included genes regulating translation, and groups 7 and 9 were enriched in genes that promote cell division (Figure 2A). The complete list of enriched GO terms is included Sup. Table 4. Seven of the 10 groups showed a general trend for reduction in transcription during differentiation to GC neurons, whereas genes in groups 2, 5, and 6 that include GO terms related to neurogenesis were transcribed at higher levels in differentiating and/or mature GC neurons compared to cycling GC progenitors.

We next examined the cilia and centrosome related GO terms associated with each gene expression group (Figure 2C). No positive regulator of pre-mitotic cilia deconstruction was revealed in the groups with increased expression in differentiating and mature GCs (groups 2 and 5). We noticed, however, that group 1, which had decreased expression in differentiating and mature GCs relative to G1 progenitor GCs, included cilia GO terms important for cilia maintenance: “ciliary transition zone”, “intraciliary transport”, “intraciliary retrograde transport", and “intraciliary transport particle B”. The transition zone at the base of the cilium creates and maintains the permeability barrier that allows for specialization of ciliary membrane and cytoplasmic content (Reiter et al., 2012). Intraciliary transport is accomplished by the IFT (intraflagellar transport) complex, which contributes to cilia assembly and maintenance by transporting protein cargo to and within the cilium (Kozminski et al., 1993; Rosenbaum and Witman, 2002; Taschner and Lorentzen, 2016). The clustering of transition zone and IFT components with genes whose expression drops off upon onset of differentiation suggested that cilia deconstruction might occur during differentiation because cilia cannot be maintained in the absence of necessary components.

This hypothesis was further supported when we examined centrosome genes. The GO terms for “centriolar satellites” and “pericentriolar material” (PCM) were enriched in group 9 which had genes whose expression was lower in differentiating and mature GCs than in cycling progenitor GCs. This was noteworthy because centriolar satellites and PCM proteins regulate ciliogenesis and promote ciliary maintenance (Werner et al., 2017).

In summary, we found no evidence for a positive regulator of cilia deconstruction in differentiating GCs. Instead, the data suggested global downregulation of cilia and centrosome genes that contribute to cilia maintenance. Although GO terms are suggestive, it was important to examine expression of individual transcripts and proteins to further test this prospective mechanism of cilia deconstruction.

### IFT transcripts and proteins decreased upon GC maturation

We began evaluating the possibility that gradual cilia deconstruction in differentiating GCs could be due to loss of ciliary maintenance by assessing the expression of IFT components. IFT proteins assemble into two distinct complexes, IFT-A and IFT-B, which regulate retrograde and anterograde transport in cilia (Rosenbaum and Witman, 2002; Taschner and Lorentzen, 2016). The IFT-A complex also regulates pre-ciliary trafficking of ciliary cargoes (Jiang et al., 2023; Mukhopadhyay et al., 2017). Loss of key proteins from either complex leads to the inability of cells to maintain cilia (Werner et al., 2017). We examined the expression pattern of IFT genes across the GC developmental lineage (Figure 3A). Only three genes coding for IFT-A components were detected above the threshold (5% of cells in the cluster). Of these, *Ift43* expression was decreased in mature neurons and *Wdr35* and *Ift22* were undetectable after the onset of differentiation. The expression pattern of IFT-B component genes showed decreased expression in differentiating and/or mature GC neurons for 10 of the 15 genes, indicating global downregulation of IFT component proteins during neural maturation.

**Figure 3.**
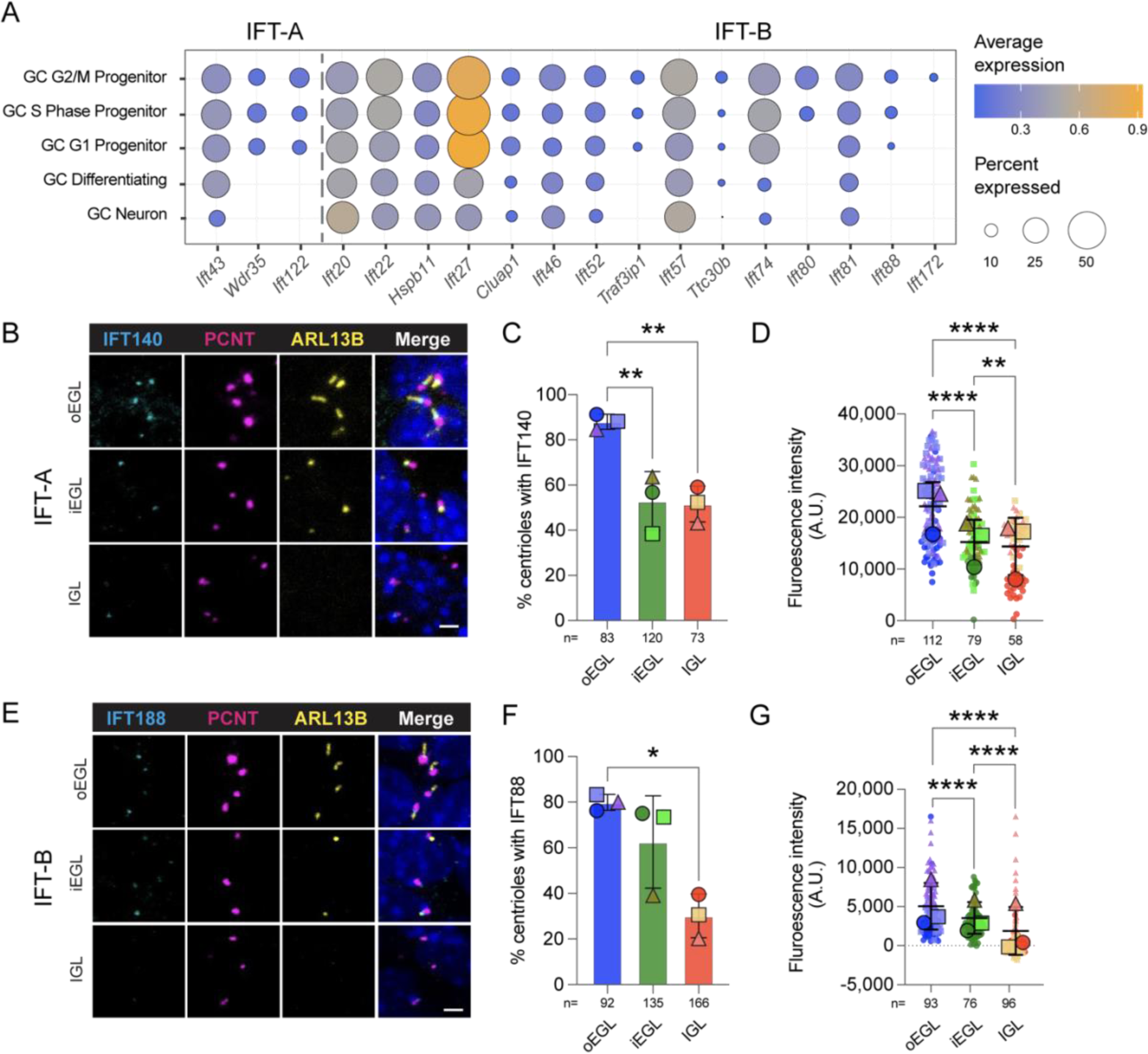
IFT transcripts and proteins decreased upon GC maturation. (A) Expression level and detection frequency of IFT genes are plotted for each GC cell cluster. Excluded IFT genes were not detected above the 5% threshold. (B, E) Sagittal sections of P7 mice cerebellum stained with antibodies to the indicated proteins and co-stained with DAPI (dark blue) were imaged by confocal fluorescence microscopy. The position of the cells in the cerebellar layers is indicated to the left of each panel. (C, D, F and G) Sections from 3 individual animals were stained and imaged. The percent of PCNT positive centrioles that also had IFT40 (C) or IFT88 (F) signal is graphed. In addition, the fluorescence intensity of each IFT40 (D) or IFT88 (G) puncta was quantified after background subtraction and the values are plotted for each layer of the developing cerebellum. Statistical analysis was preformed using multiple comparison Anova. A.U.= arbitrary units; scale bar: 1μm.

It is notable that *Ift20* expression does not diminish like other IFT components. IFT20 has been implicated in trafficking to docked centrioles at the immune synapse (Vivar et al., 2016) so it is possible expression is retained in GC neurons because the protein is similarly needed, even in the absence of the cilium. We also found additional genes related to IFT recruitment with elevated expression in differentiating and mature GC neurons (included in the expression pattern clusters 2 and 5): *Cpane1*/*Jbts17* and *Cplane2/Rsg1* code for proteins that are important for basal body recruitment of IFT-A complexes (Toriyama et al., 2016) and *Cep19* and *Rabl2* encode proteins that facilitate IFT-B injection (Kanie et al., 2017). While it is possible IFT20 and these other proteins may have non-IFT functions, we anticipate that they function at docked centrioles in adult GCs.

Transcriptional programs clearly participate in GC differentiation (Ha et al., 2019; Sato et al., 2008); however, protein levels and protein localization are more directly representative of cellular activities. If cilia deconstruction was a consequence of reductions in components necessary to maintain cilia, then decreased IFT protein levels would be expected to coincide with decreased transcription. To examine IFT protein expression and localization, we stained P7 cerebellum sections with antibodies to the IFT-A protein, IFT140, and the IFT-B protein, IFT88 (Figure 3B). In tissue sections, GC populations at different developmental stages can be distinguished based on cell location (Figure 1A) (Leto et al., 2016). Most proliferating progenitor cells populated the surface of the developing cerebellum in the outer EGL and GCs newly committed to differentiation have migrated away from the pial surface and are located in the inner EGL (Ha et al., 2019; Sato et al., 2008). As GCs progress through maturation, they migrate through the molecular layer and Purkinje cell layer and to reach the Inner Granule Layer (IGL) where neuronal maturation is completed.

In addition to staining with antibodies to IFT proteins, sections were also stained with antibodies to ARL13B to visualize primary cilia and pericentrin (PCNT), a centrosome marker. In the tissue sections, IFT88 and IFT140 both localized adjacent to centrosomes (Figure 3B, 3E). To assess changes in IFT proteins we quantified the frequency of IFT protein localization adjacent to the centrosome and the fluorescence intensity. The percentage of PCNT positive centrosomes co-staining with IFT140 decreased significantly after the onset of differentiation (Figure 3C). The percentage of centrosomes with adjacent IFT88 puncta in the outer and inner EGL was not statistically different, however, in the IGL where cilia were sparse, IFT88 detection was decreased (Figure 3E). Although it was more intense than the IFT proteins, we noticed that the frequency and intensity of PCNT seemed to diminish in the IGL. When we examined the total fluorescence intensity of the IFT proteins, we found that IFT140 intensity decreased significantly between the outer EGL and the inner EGL and remained low in the IGL (Figure 3D) and IFT88 intensity was similar in the outer and inner EGL but was barely detectable above background levels in the IGL (Figure 3F). Taken together, these data indicate that the transcriptional changes in IFT lead to reduced centrosome recruitment of the tested IFT proteins as GCs differentiate and migrate.

### Pericentriolar material (PCM) transcripts and proteins are reduced during GC differentiation

To investigate the possibility that reductions in PCM proteins during GC neural maturation also contributed to cilia deconstruction, we examined their expression patterns. As expected, based on the gene expression clustering analysis, most genes (19 out of 25 that were detected by scRNA-seq) were expressed at lower levels in differentiating and/or mature GC neurons than in cycling or G1 progenitors (Figure 4A). Specifically, we found that the gene coding for pericentrin (PCNT), *Pcnt,* was detected in fewer GCs as neurogenesis progressed. As illustrated in Figure 4B, PCNT is a PCM spoke protein that determines the circumference of the PCM (Lawo et al., 2012; Mennella et al., 2012; Pihan, 2013). PCNT was also the centrosome marker used in Figure 3 that showed decreased staining in the IGL when used as a centrosome marker in the IFT analysis. When we examined PCNT levels in immunostained sections of developing cerebellum more closely, we found that although PCNT was detectable at centrosomes in all layers, the amount of protein diminished during neural maturation (Figures 4C,D). To quantify the reductions, we measured both the total area of the centrosome signal and the fluorescence intensity within each layer of the cerebellum. We found that both were decreased in the inner EGL and then further decreased in the IGL (Figure 4E,F). Antibody accessibility is similar throughout the tissue section as evidence by PCNT fluorescence intensity in SOX9+ glia in the Purkinje layer and IGL that remained high (Farmer et al., 2016; Sun et al., 2017; Vong et al., 2015) (Sup. Figure 2). We conclude that both pericentrin transcript and protein levels diminished as GC neuronal maturation progressed.

**Figure 4.**
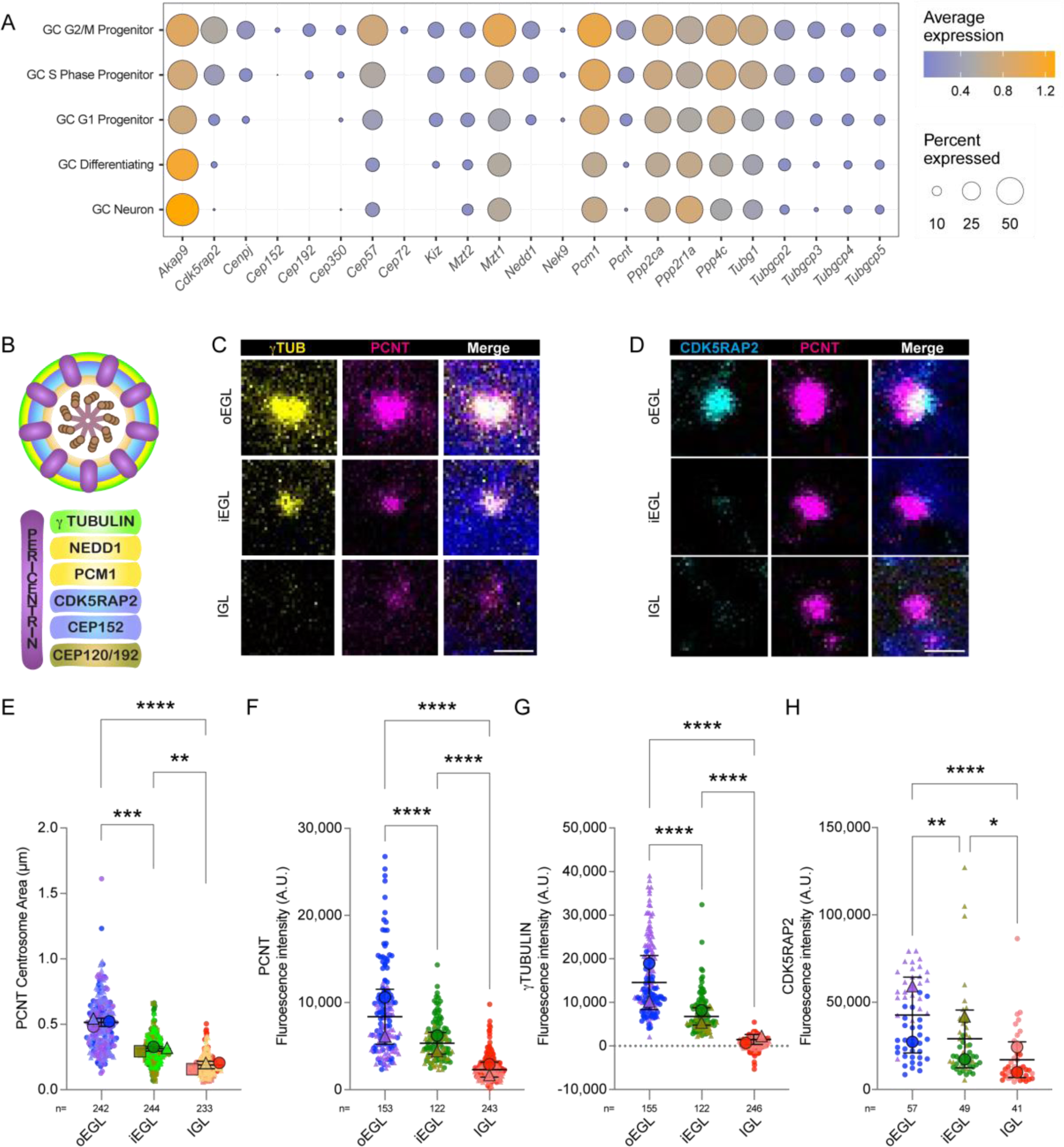
PCM transcripts and proteins are reduced during GC differentiation. (A) The average expression level and detection frequency of the indicated PCM genes are plotted for each GC cluster. (B) A representative drawing of a centrosome cross section illustrating layers of pericentriolar material. Schematic adapted from (Pihan, 2013) (C-D) Sagittal sections of P7 mice cerebellum were stained with the indicated antibodies to PCM proteins and co-stained with DAPI (dark blue) followed by confocal microscopy. The representative images are from the outer EGL, inner EGL and IGL as indicated. (E) The area of each PCNT puncta was measured and is plotted by cerebellum layer from three sections from each of 2-3 individual animals. The average measurements from each animal were superimposed as large symbols. (F-H) The fluorescent intensity of PCNT (F), γtubulin (G) and CDK5RAP2 (H) puncta in the indicated layers are plotted. Statistical analysis was preformed using multiple comparison Anova. A.U.= arbitrary units; scale bar: 1 μm.

The gene expression analysis also revealed that multiple other pericentriolar components were detected at lower levels and/or with decreased frequency in the differentiating and mature GCs. These included *Tubg1* that codes for γ-tubulin, several genes coding for members of the γ-tubulin ring complex (γ-TURC) (Lin et al., 2015), and others coding for *Nedd1 and Cdk5rap2* that are required for γ-TURC targeting and microtubule nucleation (Fong et al., 2008; Luders et al., 2006) (Figure 4A). The γ-tubulin in coordination with γ-TURC nucleates microtubules to create the microtubule organizing center for the centrosome and forms the outer layer of the PCM (Figure 4B) (Lawo et al., 2012). We therefore assessed protein levels for two of these pericentriolar proteins by staining P7 cerebellum sections with antibodies to γ-tubulin or CDK5RAP2 (Figure 4C and D). In the EGL, both γ-tubulin and CDK5RAP2 co-localized with PCNT. However, the measured fluorescence intensity of γ-tubulin and CDK5RAP2 decreased between the outer and inner EGL and was almost non-detectable in the IGL (Figure 4G and H). Thus, like the transcripts, γ-tubulin and associated proteins were largely absent from the mature GCs in the IGL.

It is interesting to note that included among the genes that were upregulated in GC neurons were the PP2A phosphatase catalytic subunit *Ppp2ca* and regulatory subunit *Ppp2r1a.* The PP2A phosphatase has been implicated in PCM dissolution (Magescas et al., 2019).

To determine whether centriole components were also depleted as PCM protein levels decreased, we imaged P7 cerebellar sections stained with antibodies to TALPID3 or CEP164 along with anti-PCNT and anti-ARL13B. TALPID3, a distal centriolar protein (Wang et al., 2016) was present in all GCs at every developmental stage (Sup. Figure 3). The distal appendage protein, CEP164 was also detected at similar levels in all GCs, irrespective of developmental stage (Sup. Figure 3). Distal appendages were visible in EM images of centrioles at all stages (Ott, Constable et al, co-submitted), and because CEP164 is recruited in the late stages of centrosome maturation (Tanos et al., 2013), we conclude that the size and composition of the PCM diminished during differentiation and maturation, while leaving the centrioles intact.

### Centriolar satellite protein expression was restricted to the outer layer of EGL

In addition to PCM proteins, the centriolar satellite GO term was identified in the expression pattern clusters that decreased in differentiating and mature GCs (Figure 2). Centriolar satellites are large protein complexes, > 65 proteins, associated with the centrosome, and are responsible for maintaining homeostasis of the centrosome, trafficking proteins to the cilium and to regulate ciliogenesis (Prosser and Pelletier, 2020). Because downregulation of centriolar satellite proteins causes reduced ciliation in cultured cells (Odabasi et al., 2019), we investigated transcript levels of centriolar satellite proteins in each GC cluster. We found that ∼70% of the genes were detected at lower levels or with lower frequency in differentiating and/or mature GC neurons compared to cycling GC progenitors or G1 GC progenitors (Figure 5A). Genes affected included *Pcm1,* the founding centriolar satellite scaffolding protein that is required for centriolar satellite structure and function (Wang et al., 2016) and *Cep131*, a gene known to be transcriptionally regulated during cerebellum development by the transcription factor ATOH1 during GC progenitor proliferation (Chang et al., 2019). Other notable genes with decreased transcription during developmental progression included *Ofd1*, *Cep290*, *Mib1*, *Sdccag8*, and *Odf2l*. To determine if downregulated transcription of centriolar satellite genes coincided with changes in protein levels, we stained cerebellum sections with antibodies to PCM1 or CEP131 along with anti-PCNT and anti-ARL13B antibodies. Centriolar satellites positive for PCM1 or CEP131 were found only in the outer most layer of the outer EGL and were completely absent from the cells in the inner EGL (Figure 5B-E, Sup. Figure 4). Along with reductions in IFT and PCM proteins, loss of centriolar satellites likely contributed to eventual deconstruction of cilia.

**Figure 5.**
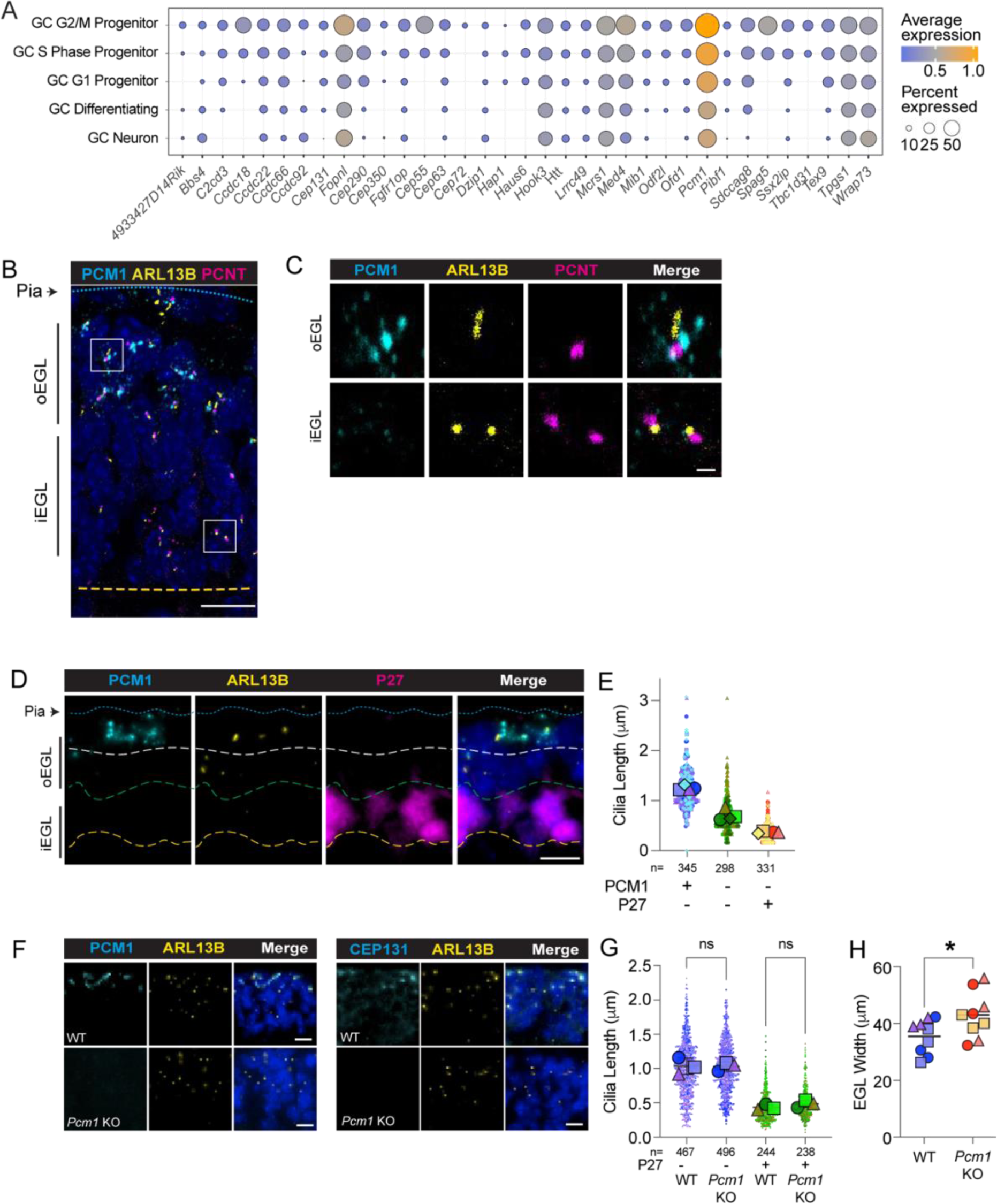
Reduction in centriolar satellite transcripts and proteins in GCs coincided with decreases in ciliary length in the inner EGL. (A) The average expression level and frequency of transcript detection for the indicated centriolar satellite genes in each GC cluster was plotted. (B-C) Sagittal sections of P9 mice cerebellum were stained with antibodies to PCM1 (centriolar satellite), ARL13B (cilium) and PCNT (PCM) and co-stained with DAPI (dark blue). The indicated regions of the outer EGL and inner EGL in B are magnified in C. (D) Sagittal sections of P7 cerebellum were stained with the indicated antibodies. P27^KIP^ is a marker of GC differentiation. The blue dashed line indicates the pial surface, the white line is the lower boundary of PCM1 positive cells, the green line is the upper boundary of the P27^KIP^ positive cells, and the yellow dashed line indicates the inside edge of EGL. (E) Cilia lengths were measured in three sections from each of three animals. Lengths were plotted based on the expression of PCM1 and P27^KIP^ and the average cilium length for each animal is indicated as a large shape. (F) Sagittal sections of P8 cerebellum from WT or *Pcm1* KO littermates stained with the indicated antibodies were imaged by widefield microscopy. (G) Cilia lengths were measured in three sections from each of three animals per group. Lengths were plotted based on the expression of P27^KIP^ and the average cilium length for each animal is indicated as a large shape. Analyzed using multiple comparison Anova. (H) Width of EGL was measured in 5 different places on 3 images each from 3 animals. Each point represents the average of 5 measurements for each section. In all merge image panels nuclei stained with DAPI are blue. Scale bar: 10 μm, zoom 1 μm.

Because we noticed that PCM1 was largely in the outermost cells in the EGL we also stained cerebellar sections with antibodies to PCM1, ARL13B and the marker of GC differentiation, P27^KIP1^, to determine if PCM1 was present in some or all P27^KIP1^ negative outer EGL progenitors. We found two layers of P27^KIP1^ negative cells in the outer EGL: an outer layer that expressed PCM1 and an intermediate layer without PCM1 signal. To investigate whether the absence of PCM1 affected cilia, we compared cilia length across the EGL. We found that cilia were shorter in the PCM1 negative progenitors than in the progenitors with centriolar satellites in the outer EGL (Figure 5E). In addition, the cilia in the P27^KIP1^ positive cells of the inner EGL were even smaller. We conclude that loss of centriolar satellites correlated with early decreases in cilia length.

To investigate whether centriolar satellites were essential for cilia formation in the EGL we sought to assess cilia formation and disassembly during GC neurogenesis in the absence of PCM1. We obtained *Pcm1* knockout mice (Monroe et al., 2020) and confirmed loss of PCM1 by immunostaining (Figure 5F). We also stained the *Pcm1^−/−^* tissue with antibodies to CEP131 and found no evidence for centriolar satellite formation in the knockout tissue (Figure 5F). Loss of PCM1 has been shown to cause age dependent cilia abnormalities in a subset of brain regions (Monroe et al., 2020). In developing cerebellar tissue, however, we found that GC progenitors still formed ARL13B positive cilia in the absence of PCM1 and centriolar satellites. Importantly, there was no difference in cilia length between *Pcm1^−/−^*pups and wild type litter mates at any stage of GC differentiation suggesting the developmental program of deconstruction were not affected in these animals. The loss of centriolar satellites did appear to effect cerebellar development: the depth of the EGL was slightly but significantly increased in the *Pcm1^−/−^* animals compared to the WT litter mates (Figure 5G). Two of the three *Pcm1^−/−^* pups were also slightly smaller than their WT littermates. Since the EGL layer decreases in thickness as the mouse ages, the increase in thickness could be attributed to a developmental delay in these animals. While systemic loss of PCM1 in a knockout animal may not be equivalent to the acute loss of PCM1 in differentiating GCs, these experiments suggest that cilia deconstruction is not the result of loss of a single component, but rather a consequence of simultaneously diminishment of several essential components in cilia and centrosome maintenance.

### SHH signaling downregulation during GC differentiation coincided with ciliary loss

We hypothesized that synchronized downregulation of diverse cilia components could have been orchestrated by the dramatic transcriptional reprograming that occurred at the onset of differentiation. In progenitor cells, SHH promotes proliferation by activating transcription factors including *Cyclin D1* and *Myc-N*, which directly drive the cell cycle (Pak and Segal, 2016; Zhao et al., 2002). Upon onset of differentiation, loss of SHH pathway activity is evident in the scRNA-seq data: expression of genes directly regulated by SHH pathway transcription factors are all reduced in differentiating and mature GCs. These included direct targets such as *Ccnd1, Gli1, Gli2, Hhip1, Mycn, Ptch1, Ptch2* and *Sfrp1* (Figure 6A, left) (Hui and Angers, 2011; Zhao et al., 2002). Genes of several other proteins that regulate SHH signaling (such as *Boc*, *Smo*) also show downregulation in differentiated vs undifferentiated GCs (Figure 6A, right). Genes coding for proteins that are negative regulators of the SHH pathway had increased expression during neurogenesis (Figure 6A, right). Several of these genes were included in the expression pattern clusters that had higher expression levels post-differentiation (clusters 2, 5 and 6 from Figure 2), such as *Tulp3* (Norman et al., 2009), *Ankmy2* (Somatilaka et al., 2020), *Sufu* (Kogerman et al., 1999), PKA regulatory subunits *Prkaca/Prkacb* (Hammerschmidt et al., 1996) and cilia localized adenyl cyclases (*Adcy3*) (Somatilaka et al., 2020; Vuolo et al., 2015). We further examined SHH pathway associated genes to determine how expression changed through GC neurogenesis. Several of these genes also had decreased expression levels or detection frequency (Figure 6B). In addition, expression of *Atoh1,* the transcription factor responsible for the proliferation and maintenance of GC precursors (Flora et al., 2009), was undetectable upon GC differentiation. ATOH1 maintains cilia and SHH responsiveness in GC progenitors partly by promoting expression of the centriolar satellite protein CEP131 (Chang et al., 2019) and by synergizing with GLI2 in activating GLI target genes (Yin et al., 2019). We conclude that the global transcriptional reprogramming that impacts expression of essential cilia and centrosome genes coincides with the loss of ATOH1 during differentiation and decrease in expression of SHH pathway components.

**Figure 6.**
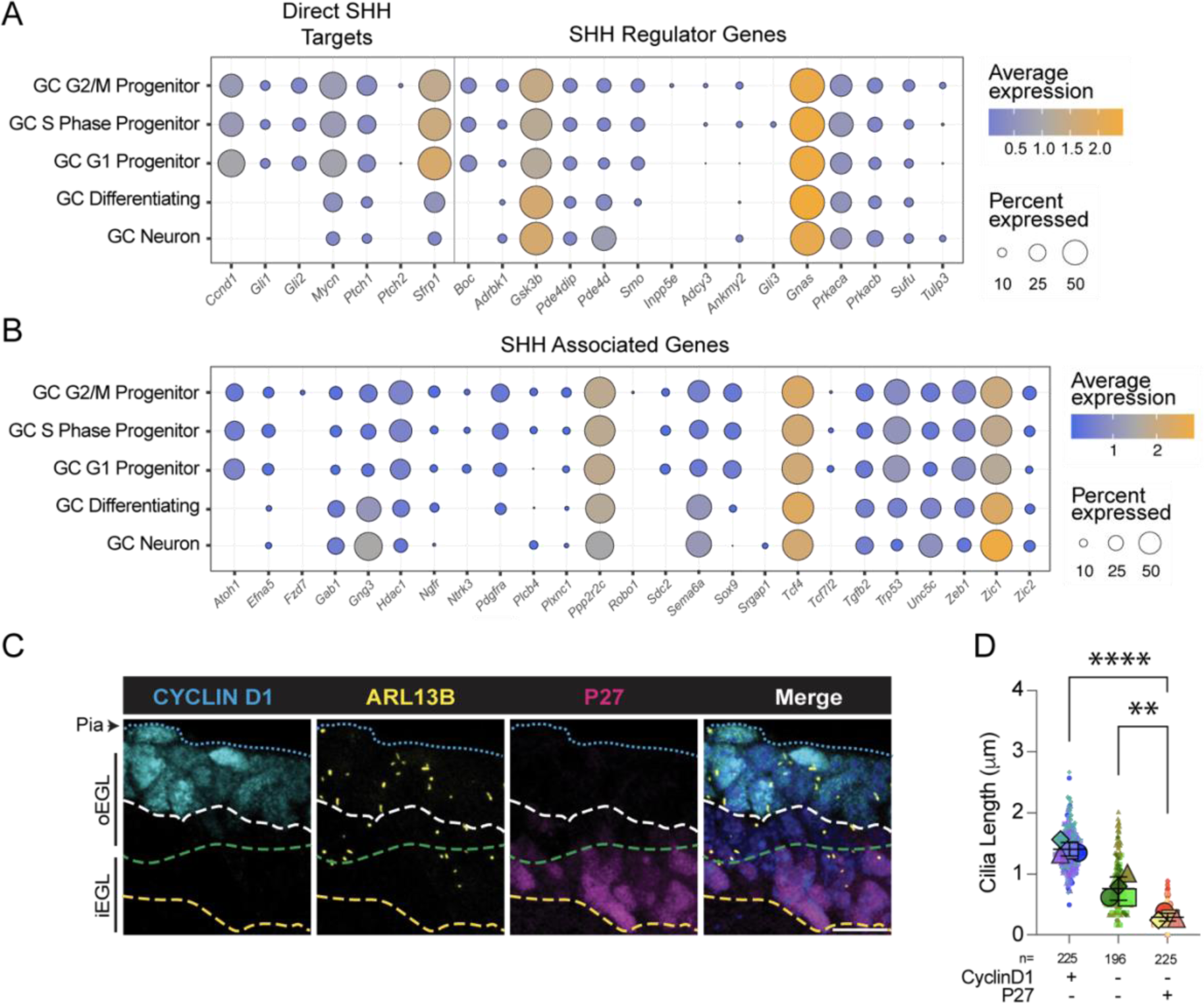
SHH signaling downregulation during GC differentiation coincided with ciliary loss. (A-B) Transcription of genes that are directly controlled by Shh pathway activity (A, left), genes coding for proteins that impact the SHH signaling cascade (A, right) and genes related to the SHH pathway (B) are graphed for each GC cluster. Transcript detection frequency is indicated by the size of each dot and the dot color represents the average expression level. (C) Sagittal sections of P7 mice cerebellum were stained with antibodies to the SHH pathway transcriptional target CYCLIN D1, the ciliary marker protein ARL13B and the GC differentiation marker P27^KIP^ along with co-staining with DAPI (blue) prior to imaging by widefield microscopy. The blue dashed line indicates the Pial surface, the white line is the lower boundary of CYCLIN D1 positive cells, the green line is the upper boundary of the P27^KIP^ positive cells, and the yellow dashed line indicates the inside edge of EGL. (D) Cilia lengths were measured in three sections from each of three animals. Lengths were plotted based on the expression of CYCLIN D1 and P27^KIP^ and the average cilium length for each animal is indicated as a large shape. Statistical analysis was preformed using multiple comparison Anova. Scale bar: 10 μm, zoom 1 μm.

In addition to reprogramming GC transcription, termination of SHH signaling may also globally reduce translation. A direct target gene of SHH is *Mycn*, a master transcriptional regulator of ribosome biogenesis (van Riggelen et al., 2010). Both the frequency and level of *Mycn* expression decreased during the final stages of neurogenesis (Figure 6A). In addition, expression pattern clusters 1 and 4, which included 37% of the top 5% of transcripts and had decreased expression in differentiating and mature GCs (Figure 2A), included many significant GO terms related to protein biogenesis. The global changes in transcription also accomplished significant reductions in translation.

To more directly evaluate the relationship between ciliation state and SHH signaling activity, we stained P7 cerebellar sections with antibodies to ARL13B, P27^KIP1^ and the direct target of SHH regulation, Cyclin D1 (encoded by the gene *Ccnd1*) (Kenney and Rowitch, 2000; Oliver et al., 2003). As expected, Cyclin D1 was present in the ciliated GC progenitors of the outer EGL (Figure 6C). We noticed that similar to the intermediate layer in sections stained with PCM1 and P27^KIP1^ (Figure 5D), there was a layer of cells below the Cyclin D1 positive progenitors that stained negative for both Cyclin D1 and P27^KIP1^. High Cyclin D1 levels during the cell cycle drive G1-S progression in GC progenitors in the absence of SHH signaling in daughter cells (Ho et al., 2020; Min et al., 2020). Therefore, the layer of cells lacking both Cyclin D1 and P27^KIP1^ were likely completing a final cycle prior to differentiation triggered by G1-S progression due to Cyclin D1 levels from the previous cell cycle. We next measured cilia length to try to assess whether changes in primary cilia correlated with the termination of SHH signaling or the onset of differentiation. We found that the GCs negative for both Cyclin D1 and P27^KIP1^ had shorter cilia than the Cyclin D1 positive progenitors (Figure 6D), and the cilia in the cells that were P27^KIP1^ positive were even shorter (Figure 6D). Together the data indicate that cessation of SHH signaling coincided with reduced cilia length in GC progenitor cells. We conclude that along with the observed changes in both transcript and protein levels of essential centrosome and cilia components, the analysis of SHH pathway targets provided evidence for global changes that reduce the ability of GCs to maintain cilia.

### Capping proteins prevented cilia elongation from docked centrioles

Docking of mother centrioles to the plasma membrane primes the formation of cilia from the cell surface (Francis et al., 2011), and it is unusual that centrioles dock in GC neurons without forming cilia (Ott, Constable et al, co-submitted). Prior to ciliogenesis, the CEP97 and CP110 capping complex needs to be removed from the distal mother centriole (Chen et al., 2002; Spektor et al., 2007). We investigated the possibility that the capping complex was instrumental in preventing ciliary regrowth from docked mother centrioles during GC neuronal maturation. We immunostained P8 cerebellar tissue for the cap protein CEP97 along with the ciliary protein ARL13B and the pericentriolar protein PCNT (Figure 7A). The capping protein always associates with mature daughter centrioles (O’Toole and Dutcher, 2014; Spektor et al., 2007), so every centrosome had at least one CEP97 puncta. Throughout the tissue there were six different configurations of cilia, centrosomes and caps as shown in Figure 7B. As expected, ciliated centrioles did not have CEP97 signal associated with the basal body and progenitors with duplicating centrioles had two or three CEP97 puncta. We classified centrioles with cilia and large bright PCNT staining separately because those were typically located in the Purkinje cell layer and were likely ciliated glia or Purkinje neurons (Sup. Figure 2). The non-ciliated centrioles were classified as having either one or two CEP97 puncta. In several cases only a single PCNT puncta was present with two CEP97 puncta, which we believe happened when the mother and daughter centriole were too close to resolve.

**Figure 7.**
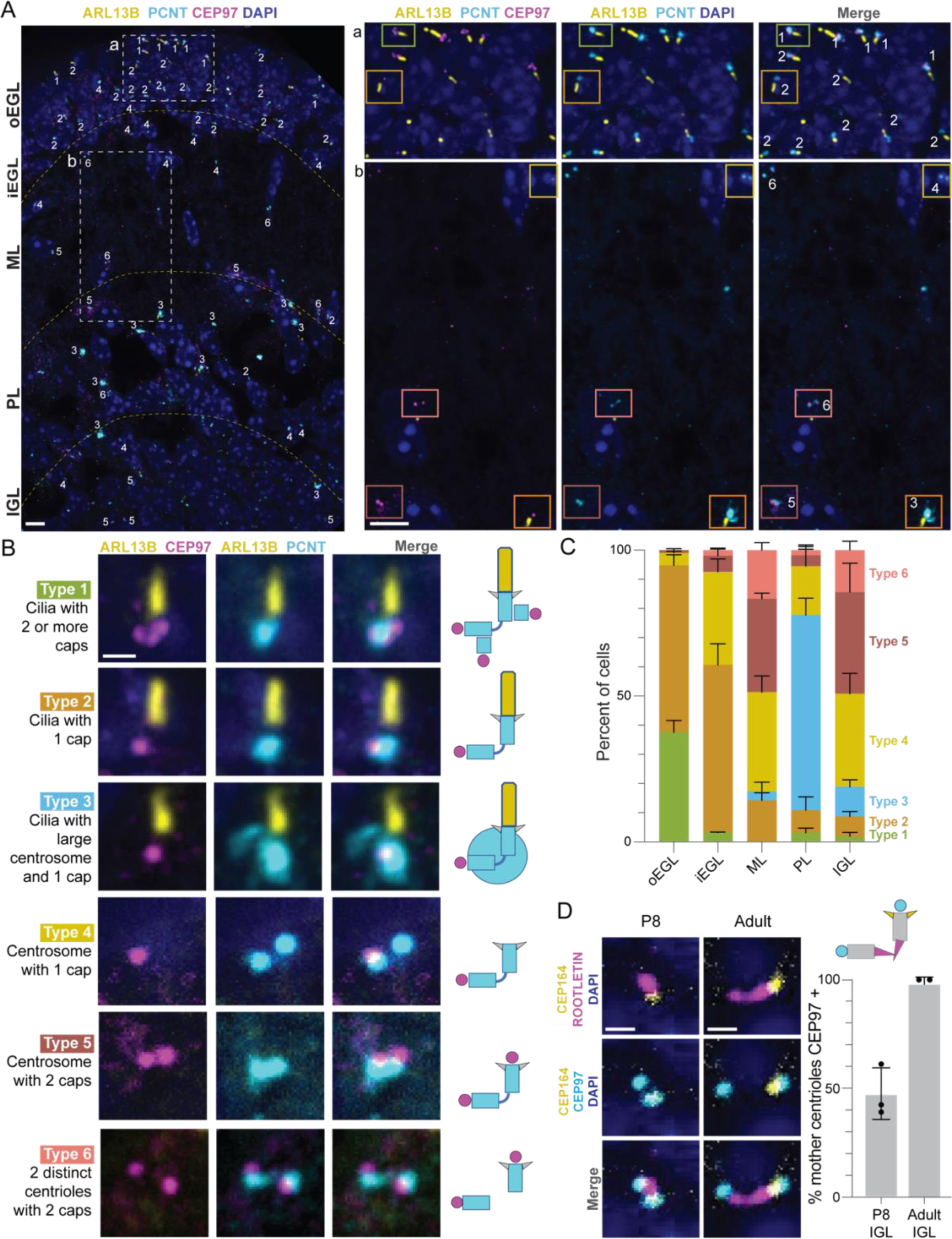
Centriole capping in mature GC neurons prevented regrowth of primary cilia. (A) Sagittal sections of P8 mice cerebellum sections stained with indicated antibodies and co-stained with DAPI were imaged using spinning disk confocal microscopy. Dotted lines indicate boundaries between cerebellar layers. The boxed regions are magnified and shown as channel subsets to the left. Centrosomes were classified based on the presence or absence of a cilium, the area of the PCNT centrosome puncta and the number of adjacent CEP97 puncta. (B) Representative images and an illustration of each class of centriole are shown for each class. Each image corresponds to the square in A with coordinating color. (C) The distribution of centrosome type is graphed for each layer. 210-327 centrioles per animal were classified from 3 animals (D) Sagittal sections from adult and P8 cerebellum were stained with antibodies against the distal appendage protein CEP164, distal centriole associated rootletin, the capping protein CEP97 and counter-stained with DAPI before imaging as in (A). The percentage of centrosomes with two CEP97 puncta is graphed. 36-145 mother centrioles were assessed from each of 3 mice. Adult mice were aged P39/P50/P160.

To evaluate changes in CEP97 binding to mother centrioles during GC development, we quantified the distribution of the centriole classes across each layer (Figure 7C). We noted that ∼45% of the outer EGL GCs had cilia and either two or three CEP97 puncta. A similar percentage of ciliated GCs in the outer EGL and ∼60% GCs in inner EGL had a single CEP97 puncta indicating they were G1/G0 phase cells. In the inner EGL, centrioles without cilia typically had a single CEP97 puncta. The fraction of cells with no cilia and two capped centrioles increased as differentiation progressed. In the IGL, centrioles with two CEP97 puncta were more frequent than cells with an uncapped centriole, suggesting that capping of GC mother centrioles occurs as differentiation progresses. To determine whether the docked mother centrioles in mature GC neurons remained capped to prevent ciliogenesis, we sectioned and stained both P8 and adult cerebellar tissue with antibodies to CEP97 and two markers: the distal appendage protein CEP164 and the protein rootletin, which associates with the distal end of the centriole. Like the staining in Figure 7C, about half of the centrosomes in the P8 IGL had two CEP97 puncta associated with each centrosome. In the adult tissue, however, 100% of the cells had two Cep97 puncta (Figure 7D), indicating that both the mother and daughter centrioles were capped which prevented ciliogenesis in GC neurons. We conclude that although depletion of ciliary components led to the deconstruction of cilia during neuronal maturation, an additional mechanism, centriole capping, prevented cilia from reforming, despite the anchoring of the mother centriole to the plasma membrane.

## DISCUSSION

This study and the accompanying manuscript reveal structural and molecular changes in cilia and centrosomes that occur *in vivo* during GC neurogenesis. Correlation of insights from analysis of existing electron microscopy and scRNA-seq datasets, in combination with immunocytochemistry, enabled us to define a cilia disassembly pathway in differentiating GC neurons distinct from pre-mitotic cilia disassembly. Specifically, we show that cilia deconstruction in GC neurons occurs because multiple components required to maintain primary cilia are downregulated. In addition, we discovered global changes in transcription and translation, initiated at least in part by downregulation of SHH signaling that coincided with reductions in cilia length. The process of cilia deconstruction is not only set in motion by the shift in transcriptional program, but also shows a lack of a molecular activation trigger compared to pre-mitotic cilia resorption (Pugacheva et al., 2007; Tucker et al., 1979). The gradual decrease in factors required for cilia maintenance could dictate the wide spatiotemporal distribution of the ciliary deconstruction intermediates during GC neuronal differentiation (Ott, Constable et al. co-submitted). Moreover, cilia regrowth was prevented after cilia deconstruction by recruitment of CEP97, a component of the centriolar cap that prevents cilia extension.

To our knowledge, GC cilia deconstruction is the first example where withdrawal of cilia maintenance is the proposed mechanism for programmed cilia disassembly. Organelle maintenance has been defined as “all processes in a cell that prevent a given property from deteriorating” (Werner et al., 2017). For cilia and centrosomes, maintenance has largely been considered relevant to organelle stability (Izquierdo et al., 2014; Pimenta-Marques et al., 2016) and aging related phenotypic changes (Cornils et al., 2016). The requirement for IFT has been demonstrated in multiple experiments. *Chlamydomonas* flagella shorten upon inhibition of anterograde IFT in temperature sensitive kinesin-II mutants (Marshall et al., 2005; Marshall and Rosenbaum, 2001). Acute inhibition of kinesin-II also results in cilia loss in NIH 3T3 cells (Engelke et al., 2019). Both centriolar satellites and the PCM are thought to be essential for cilia maintenance because they facilitate recruitment and possibly assembly of cilia components. The peri-basal body recruitment of IFT has been shown to be regulated by diffusion and capture by the basal body rather than by microtubular transport (Hibbard et al., 2021). For example, loss of the distal appendage localized kinase TTBK2 resulted in a reduction of IFT88 recruitment to the base of the cilium (Nguyen and Goetz, 2022). Satellite proteins such as PCM1 also regulate IFT levels in and around cilia (Hall et al., 2023). In GCs the loss of PCM and centriolar satellite proteins could undermine cilia maintenance in part because they are needed for IFT recruitment.

Capping of docked mother centrioles has only been reported in cytotoxic T cells. In that case, the CP110/CEP97 complex remains with the mother centriole upon membrane docking (Stinchcombe et al., 2015). We show that in differentiating GC neurons, capping occurs following cilia loss. Although the temporal relationship between capping and docking of mother centrioles in GC neurons is presently not precisely known, the emergence of CEP97 capping appeared to correlate with centriole docking observed by electron microscopy (Ott, Constable et al. submitted)(Del Cerro and Snider, 1969). Both docking and capping were incomplete in the IGL during postnatal development, while in adult tissue almost every centriole had docked centrioles by electron microscopy and capped centrioles by immunofluorescent microscopy. In cultured RPE cells the absence of PCM1 caused CP110 and CEP97 to be retained at the centrosome and the cells failed to ciliate (Hall et al., 2023). It is possible that the depletion of centriolar satellite proteins also facilitates centriole capping following cilia loss.

Medulloblastomas are cerebellar granule cell tumors. GC tumor cells in both the SHH- and WNT-subtypes of medulloblastoma are ciliated (Han et al., 2009; Youn et al., 2022). Development of SHH-subtype medulloblastoma is preceded by formation of persistent proliferative nests of GCs (Oliver et al., 2005; Shimada et al., 2018). Additional therapeutic targets are needed because the use of Smoothened inhibitors, while initially promising (Kool et al., 2014), is restricted in young patients because it can cause skeletal growth retardation, premature growth plate closure, and drug resistance (Kieran et al., 2017; Robinson et al., 2017; Zhao et al., 2015). It is not yet clear whether the ciliated cells in tumors failed to deconstruct cilia, failed to cap cilia, or regrew cilia by reversal of cilia deconstruction and capping. Future studies will be required to determine how transcription and translation of IFT, PCM and centriolar satellite components differ in ciliated GCs of the proliferative progenitor nests and in tumors. This knowledge could facilitate targeted therapeutic strategies that could be explored as a novel approach to treat ciliated medulloblastoma subtypes.

## Acknowledgements

This project was funded by the National Institutes of Health (1R35GM144136 to S.M and 1S10OD028630 to Microscopy Core Facility in UT Southwestern), an Alex Lemonade stand Foundation A-Award to S.M., and the Howard Hughes Medical Institute. The authors would like to acknowledge the Quantitative Light Microscopy Core, a Shared Resource of the Harold C. Simmons Cancer Center in UT Southwestern, supported in part by an NCI Cancer Center Support Grant, 1P30 CA142543-01. The content is solely the responsibility of the authors and does not necessarily represent the official views of the National Institutes of Health. Figure 1 and the graphical abstract contain drawings created in part with BioRender.com. We thank molecular pathology and mouse animal care facility in UT Southwestern. We thank Dr. Gregory Pazour for his kind gift of antibodies. We also thank Andy Moore for discussions about figures and Christina Gladkova, Lauren Porter, Lorena Bendetti, Chris Obara, and Cayla Jewett for helpful discussions and comments on the manuscript.

## Author contributions

S.C., C.M.O. and S.M. conceived the project. A. Lemire performed the transcriptomic investigations, which were designed and analyzed together with C.M.O. S.C. and K.W performed immunofluorescence experiments and quantified the results with help from A. Lim. S.C., C.M.O., J. L.S. and S.M. wrote the paper with inputs from all authors.

## Materials and Methods

### scRNA-seq Clustering and analysis

scRNA-seq gene expression matrices for mouse cerebellum P5 (GSM3318005), P7 (GSM3318006) and P14 (GSM3318007) developmental time points were imported into Seurat v4.2.1 and combined (see https://figshare.com/s/6c7884fbc44230023ebd for R notebooks). Briefly, single cells with fewer than 500 genes or greater than 10 percent mitochrondrial RNA were excluded. The resulting expression data were normalized (“LogNormalize”, scale.factor = 10000) and scaled (features = all genes). Cells were clustered by gene expression and UMAP projections using the first 20 principal components were created. Positive marker genes were identified if expressed in a minimum of 25% of cells within a cluster with a minimum average log2 fold-change threshold of 0.25 (only.pos = T, min.pct = 0.25, logfc.threshold = 0.25).

### Gene expression pattern clustering

The top 5% highest expressing genes within GCs were identified and combined with the curated list of genes for hierarchical clustering to identify patterns of gene expression within each GC cluster. The curated list of cilia and centrosome genes was generated based on the literature. In Sup. Table 2 general references are included in the first tab and references for select individual genes are included adjacent to the gene name.

Using normalized scaled gene expression values from GC cell clusters, we calculated the mean expression within each GC cluster, then calculated the global mean expression per gene across all GCs and selected the top 5% (see https://figshare.com/s/6c7884fbc44230023ebd for R notebooks) for subsequent analysis. Hierarchical clustering of gene expression was performed on GCs and the resulting dendrograms were cut with varying group sizes (k values). The gene expression groups were evaluated by inspecting groupings in the dendrograms and performing GO analysis of genes within each expression group using DAVID. A cutoff of k=10 was chosen for the combined list of the top 5% of expressed genes plus the curated genes, which performed similarly to a reduced set of genes (excluding curated genes) using a cutoff of k=9.

The genes included in each expression pattern cluster are included in Sup. Tables 3 and 5. To identify gene families with similar gene expression patterns, we used the Database for Annotation, Visualization and Integrated Discovery (DAVID) (Sherman et al., 2022). The GO terms enriched in each cluster are listed in Sup. Tables 4 and 6. For analysis, we considered Biological Process (BP), Cellular Component (CC), or Molecular Function (MF) GO terms with statistical enrichment indicated by a Benjamini score < 0.005.

### Mouse handling and genotyping

All animal studies were approved in accordance with UTSW Institutional Animal Care and Use Committee regulations and were conducted in accordance with NIH guidelines for the care and use of laboratory animals. CD1 mice were purchased from Jackson Labs and maintained under standard conditions. Mice mutant for *Pcm1* were a gift from Nicholas Katsanis, Northwestern University Feinberg School of Medicine (Monroe et al., 2020). Mouse genotyping was performed as described in Monroe et al., 2020 by PCR of genomic DNA obtained from ear biopsies or toe clips.

### Mouse brain processing

Mice were procured at the appropriate age and fixed by trans-cardial perfusion using 4% paraformaldehyde (PFA) in PBS after appropriate anesthesia for their age (either isoflurane or cold exposure on ice) according to IACUC regulations. Brains were removed and further fixed in 4% PFA/PBS overnight at 4°C on a rotator, then immersed in 30% sucrose in PBS until brain sank to the bottom of the tube (∼48hrs). Brains were cut in half in the sagittal direction and embedded cut face down in cryomolds using OCT embedding media (BioTek, USA) and frozen on dry ice until solid. Blocks were stored at −80°C until sectioning on a Leica Cryostat model CM1950 at 15-30 μm thickness. Sections were stored at −20°C or −80°C until staining.

### Immunofluorescence staining and light microscopy

Sections were thawed at room temperature and OCT was removed by immersion in PBS. Sections were blocked using 3% serum (donkey) in PBS with 0.3% Triton-X 100 for 30mins. Primary antibodies were diluted in blocking solution at the appropriate dilutions and incubated overnight at room temperature in humid chamber. Primary antibodies: ARL13B (1:1000, UC Davis/NeuroMab #75-287), CDK5RAP2 (1:500, Bethyl #IHC-00063-T), CEP131 (1:500, Proteintech #25735-1-AP), CEP164 (1:500, Proteintech #22227-1-AP; 1:200, Proteintech #CL488-22227 (Fig 7D)), CEP97 (1:200, Proteintech #22050-1-AP), Cyclin D1 (1:500, NeoMarkers #RB-9041-P0), γTUB (1:500, Santa Cruz #sc-17787), P27^KIP^ (1:400, BD Biosciences) #610241), PCM1 (1:500, Bethyl #A301-149A-T), PCNT (1:500, BD Biosciences #611814), Rootletin (1:500, Merck-Millipore #ABN1686), SOX9 (1:500, Millipore #ABE571), TALPID3 (1:500, Proteintech 24421-1-AP). Rabbit polyclonal antibodies against IFT140 (1:500), IFT57 (1:500) and IFT88 (1:500) were kind gifts of Dr. Gregory Pazour, UMass Med School and required incubation with 0.05% SDS for 5 mins followed by several washes with PBS before blocking solution was applied. Sections were incubated with the indicated secondary antibodies for 2 h at room temperature. When two mouse antibodies were used, isotype specific secondary antibodies were used. To stain nuclei, Hoechst 33342 (10μg/ml) was included with the secondary antibodies or DAPI (1μg/ml) Sigma) was added to the final wash. Stained tissues were mounted using Fluromount-G (Southern Biotech) and allowed to dry overnight. Stained slides were imaged within 2-3 days and stored at 4°C (short term) or - 20°C (long term).

Images were acquired on a widefield microscope (AxioImager.Z1; ZEISS), confocal microscope (Zeiss LSM880) or a spinning disk confocal microscope (Nikon CSU-W1 SoRa). Images in the widefield microscope were acquired using a Plan Apochromat objective (40×/1.3 NA oil and 63×/1.4 NA oil) and sCMOS camera (PCO Edge; BioVision Technologies) controlled using Micro-Manager software (University of California, San Francisco) at room temperature. Images in the confocal microscope (Zeiss LSM880) were acquired using Plan Apochromat objective (63×/1.4 NA oil). Images in the spinning disk confocal microscope (Nikon CSU-W1 SoRa) were acquired using a Plan Apochromat objective (100×/1.45 NA oil), a sCMOS camera (Hamamatsu Orca-Fusion), and a Piezo z-drive for fast z-stack acquisition controlled using Nikon NIS-Elements software at room temperature. Between 10 and 30 z sections at 0.2 µm intervals were acquired.

### Image analysis

#### Cilia length and number determination

Images were analyzed using FIJI (Schindelin et al., 2012). Different cerebellum layers were identified using nuclei, and P27 markers. Cilia length was manually determined by tracing in each zone at zoom level 200-300% using the freehand draw tool, and measurement recorded using measure tool. Cilia already traced were permanently marked with draw tool to ensure unique cilia were measured when moving around the image. Zones were completed in their entirety before moving on to the next zone. The number of cilia was determined by counting the number of cilia measured. The number of cells was determined by counting the total number of nuclei found in each section as stained by DAPI.

#### Centrosome area and intensity measurements

Centrosomal area was determined on calibrated images by manually drawing around the centrosome as determined by PCNT staining and measured using the Measurements tool in FIJI. Integrated density of centrosome-associated proteins was determined by following established protocols (Bowler et al., 2019; Keller et al., 2014). Briefly, centrosome intensity was measured within a defined area of a constant size (20px, approx. 1.5mm^2^) encircling the centrosomal unit (centrosome +PCM) from the summed intensity projections of Z stacks. Intensity of the background in a near proximity of each centrosome was subtracted from the signal intensity. At least 40 centrosomes were measured for each condition/cell cycle stage.

### Graphing and statistics

All graphs and statistics were generated using Prism (GraphPad) or R. Superplots were generated by overlaying average values from each animal onto individual values, as explained in (Lord et al., 2020). Statistical significance was determined in Prism using ordinary one-way Anova using multiple comparison analysis with Turkey correction. Population mean was assessed at a 95% confidence interval and were considered significant at the following p values: 0.0332 (*), 0.0021 (**), 00002 (***), <0.0001 (****).

## Supplemental Figures

**Supplemental Figure 1.**
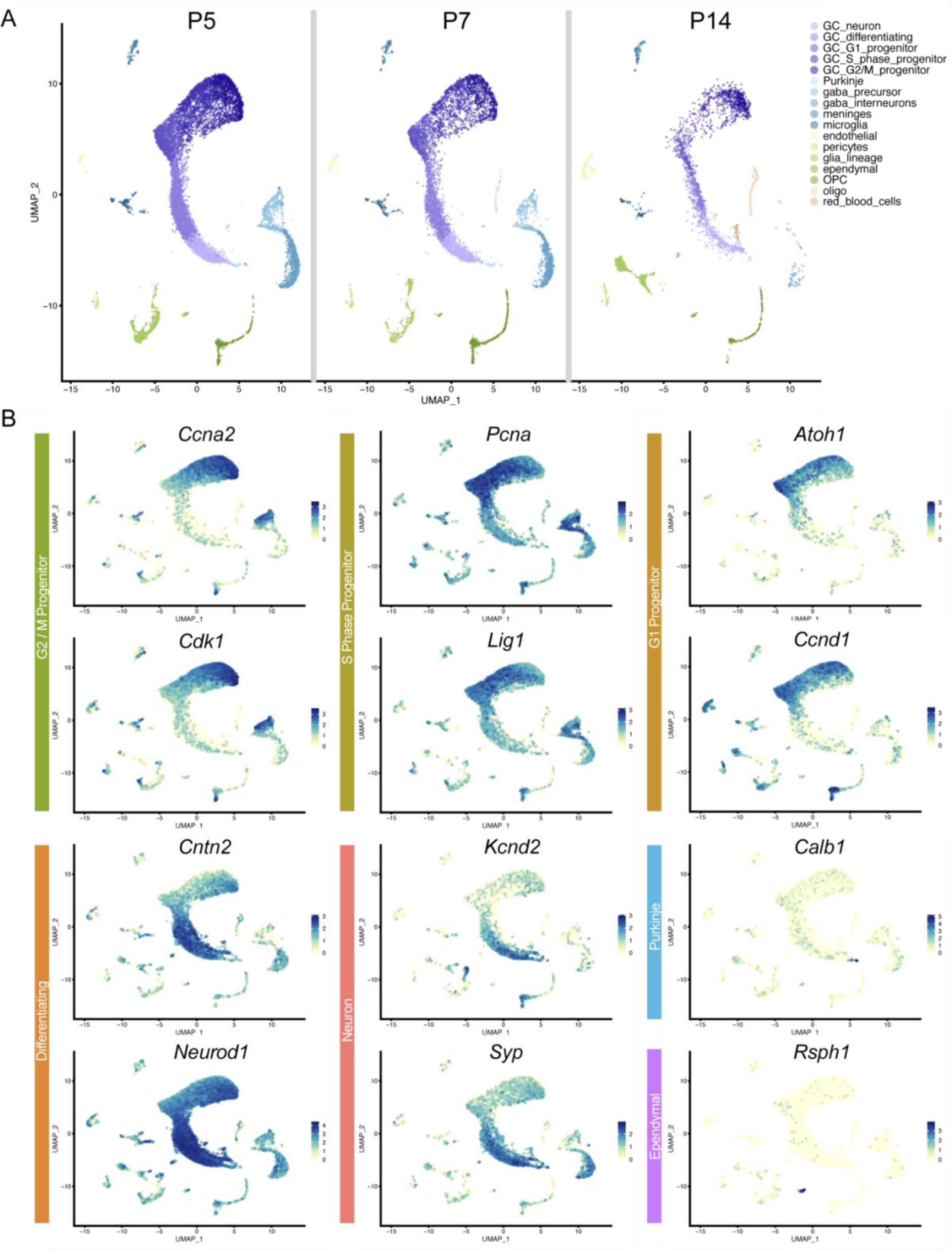
Cell cluster expression pattern distributions. (A) Cluster analysis was performed on the combined data from scRNA-seq from P5, P7 and P14 mice. Here the cluster assignments shown in Figure 1B were applied to the input data from each individual dataset. (B) The expression of the indicated genes across the combined dataset is shown. These genes are representative of genes with expression significantly enriched in the indicated cell cluster.

**Supplemental Figure 2.**
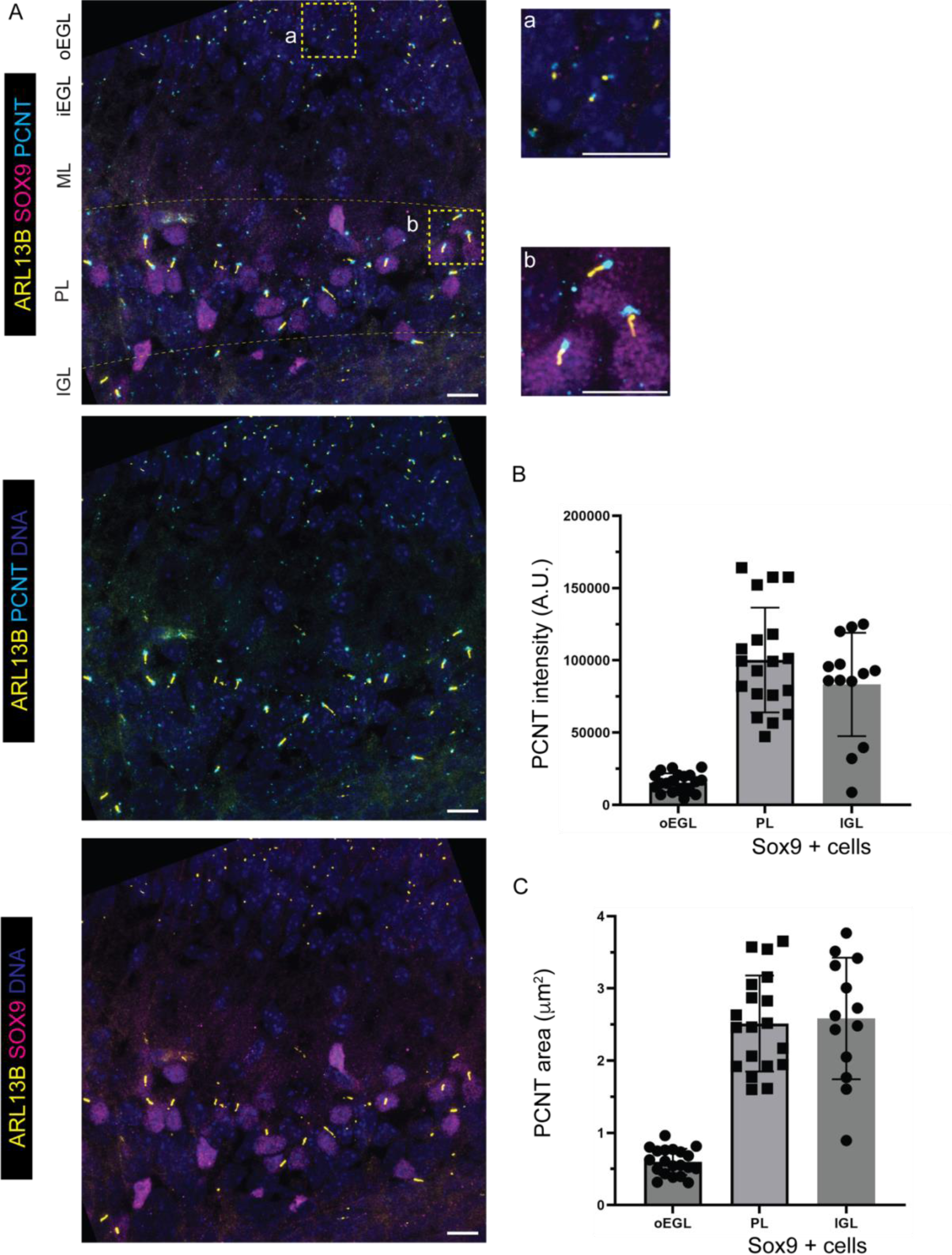
PCNT can be detected in Purkinje cells and glial cells in the PCL and IGL. (A) Sagittal sections of P8 mice cerebellum were stained with antibodies to the cilia marker ARL13B, the glial marker SOX9, the PCM marker PCNT and counterstained with DAPI. The yellow boxes indicate the location of the insets shown to the right (B and C) The area and intensity of the fluorescence intensities of PCNT signal was measured in 3 sections from 1 animal. The intensity and area are plotted for the GCs in outer EGL, and the Sox9+ glia in the PL and IGL.

**Supplemental Figure 3.**
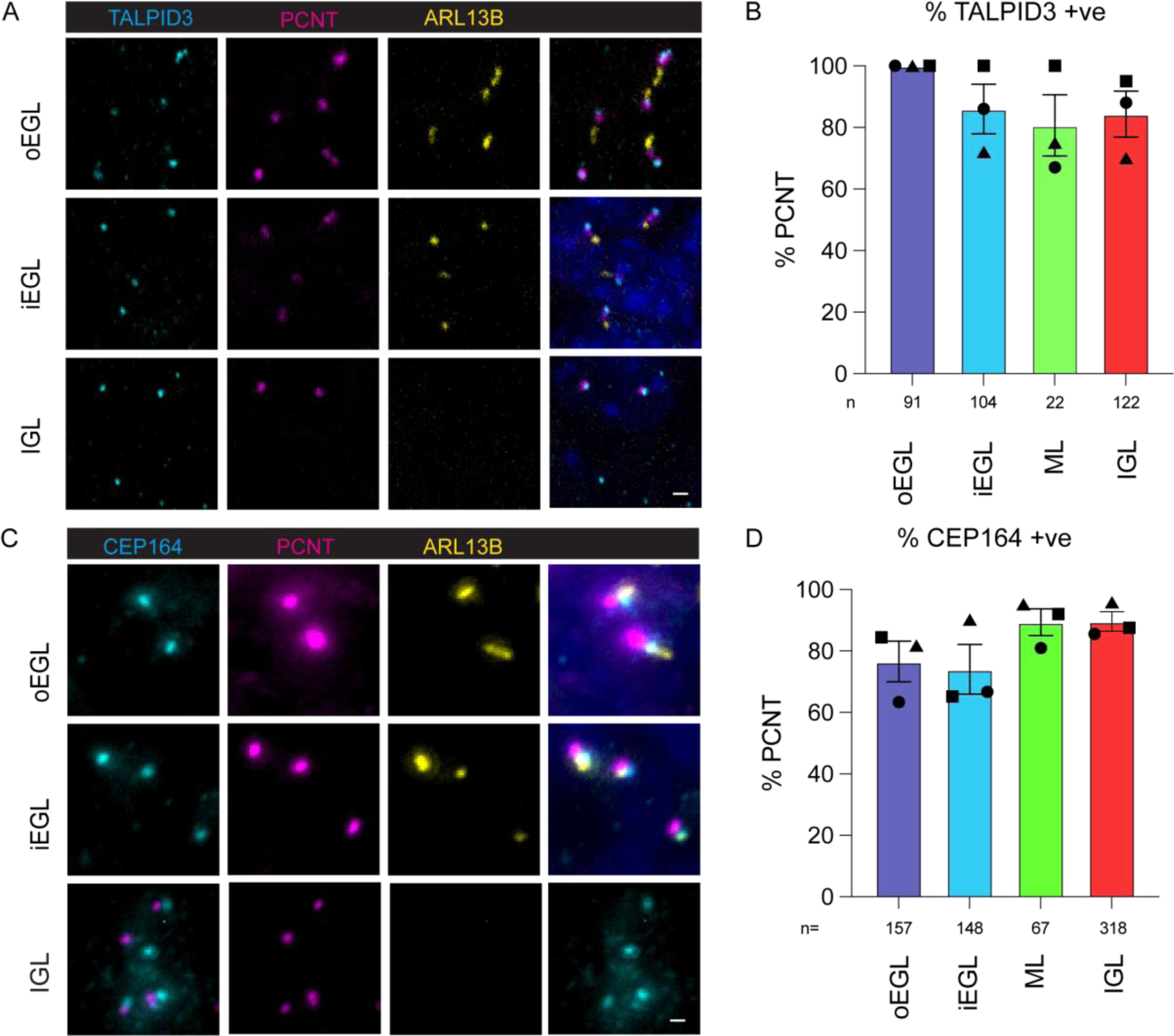
Centriolar proteins are retained during GC maturation. (A and C) Sagittal sections of P7 cerebellum were stained with antibodies to the centriole proteins CEP164 (A) or TALPID3 (C), the PCM marker PCNT, the cilia marker ARL13B, and counterstained with DAPI before imaging with confocal microscopy. Representative images from the outer and inner EGL and the IGL are shown. (B and D) The frequency of CEP164 (B) or TALPID3 (D) was adjacent to PCNT positive centrioles was measured and is plotted as a percentage of total PCNT puncta in each layer. Both the distal appendage protein, CEP164, and the centriolar protein, TALPID3, were localized with PCNT in all GCs, irrespective of maturation. Quantifications was performed from one section per animal from 3 individual animals.

**Supplemental Figure 4.**
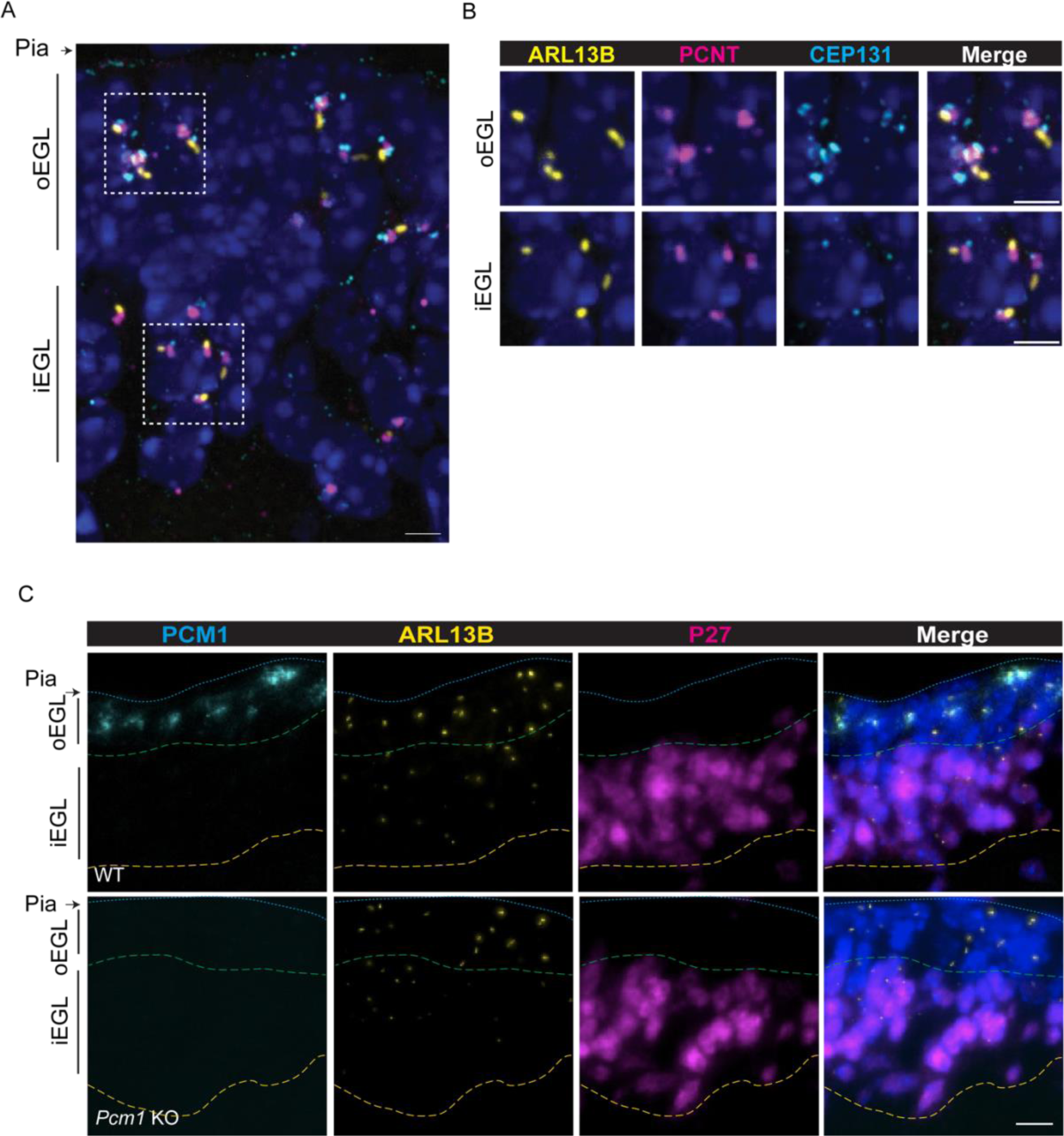
Centriolar satellites are lost during GC maturation. (A-B) Sagittal section of P7 mice cerebellum were stained with antibodies to ARL13B, PCNT, the centriolar satellite protein CEP131, and counter-stained with DAPI before imaging using confocal microscopy. CEP131 detection was decreased in differentiating GCs. The indicated regions of the outer EGL and inner EGL in A are magnified in B. Scale bar 2.5 μm. (C) Sagittal sections of P8 cerebellum from WT or *Pcm1* KO littermates were stained with antibodies to PCM1, ARL13B and P27^KIP^, and counterstained with DAPI before imaging using widefield microscopy. The blue line indicates the pia, the green line is boundary between the outer and inner EGL and the yellow line marks the beginning of the ML.

